# SUFFICIENT ACTIVITY OF THE UBIQUITIN PROTEASOME SYSTEM IN AGED MICE AND DURING RETINAL DEGENERATION SUPPORTS DHFR-BASED CONDITIONAL CONTROL OF PROTEIN ABUNDANCE IN THE RETINA

**DOI:** 10.1101/2021.04.13.438468

**Authors:** Hui Peng, Prerana Ramadurgum, DaNae R. Woodard, Steffi Daniel, Marian Renwick, Bogale Aredo, Shyamtanu Datta, Bo Chen, Rafael Ufret-Vincenty, John D. Hulleman

## Abstract

The *Escherichia coli* dihydrofolate reductase (DHFR) destabilizing domain (DD) serves as a promising approach to conditionally regulate protein abundance in a variety of tissues. In the absence of TMP, a DHFR stabilizer, the DD is degraded by the ubiquitin proteasome system (UPS). To test whether this approach could be effectively applied to a wide variety of aged and disease-related ocular mouse models, which may have a compromised UPS, we evaluated the DHFR DD system in aged mice (up to 24 mo), a light-induced retinal degeneration (LIRD) model, and two genetic models of retinal degeneration (*rd2* and *Abca4^−/−^* mice). Aged, LIRD, and *Abca4^−/−^* mice all had similar proteasomal activities and high-molecular weight ubiquitin levels compared to control mice. However, *rd2* mice displayed compromised chymotrypsin activity compared to control mice. Nonetheless, the DHFR DD was effectively degraded in all model systems, including *rd2* mice. Moreover, TMP increased DHFR DD-dependent retinal bioluminescence in all mouse models, however the fold induction was slightly, albeit significantly, lower in *Abca4^−/−^* mice. Thus, the destabilized DHFR DD-based approach allows for efficient control of protein abundance in aged mice and retinal degeneration mouse models, laying the foundation to use this strategy in a wide variety of mice for the conditional control of gene therapies to potentially treat multiple eye diseases.

## INTRODUCTION

The number of gene therapy-based approaches for the treatment of eye diseases has increased substantially over the last decade^1^. These efforts have been driven in part due to two phenomena: i) the eye is an alluring target organ for gene therapy due to its ease of accessibility and immune-privilege^2^, and ii) the FDA approval of a gene replacement therapy for the treatment of Leber’s congenital amaurosis 2 using recombinant adeno-associated virus (rAAV) to drive expression of the wild-type (WT) *RPE65* gene in retinal pigment epithelium (RPE) cells^3^. In this particular instance, constant expression of a WT copy of the *RPE65* gene is appropriate for preventing further retinal degeneration due to loss-of-function *RPE65* mutations. However, in certain instances, such as gain-of-function diseases, or scenarios wherein constitutive expression of a gene of interest may cause phenotoxicity^4^, a controllable or conditional gene therapy approach may be more desirable.

Conventional inducible gene regulation methods, such as tetracycline/doxycycline-based transactivation systems (i.e., Tet-On^5^ and Tet-Off^6^), require an extended timeframe to activate and reverse due to additional processing time of transcription and translation to reach expected protein levels, as well as prolonged deposition of the regulating molecule^7–9^. In contrast to transcriptionally-regulated conditional systems, protein-based regulation approaches such as the *Escherichia coli* dihydrofolate reductase (DHFR) destabilized domain (DD) system^10^ can directly and conditionally regulate abundance at the protein level after addition of trimethoprim (TMP)^10–12^ or a TMP-like molecule^13–15^. Such a system is especially useful when expressed transgenes are potentially toxic if expressed for prolonged periods of time, or when spatio-temporal regulation is required, such as in the case of particular stress-responsive signaling pathways which typically occur in a sinusoidal-like manner^16^.

The effectiveness of the regulation of the DHFR DD system relies on two primary processes: i) degradation of the DHFR DD fusion protein mediated through the ubiquitin proteasome system (UPS) in the absence of a stabilizer (thus determining the degree of basal expression), and ii) stabilization of the DHFR DD by TMP or TMP analogs to increase protein abundance^11, 17–22^ (this determines the fold induction). Given TMP’s broad ability to distribute across bodily tissues^23, 24^, combined with its nanomolar AC_50_ in stabilizing the DHFR-DD^14^, the latter phenomena (*in vivo* stabilization by TMP) is typically not a concern. However, previous reports have indicated that aging^25–28^ and retinal degeneration^29–34^ may correlate with compromised targeting of substrates to the proteasome or compromised proteasome activity itself, which, if true, could lead to higher levels of basal expression of the DHFR DD, and potentially compromise researchers’ ability to achieve a proper fold induction after stabilizer addition.

Previously we demonstrated that the DHFR DD system can regulate the expression of yellow fluorescent protein or firefly luciferase as a proof-of-concept in the retina of young and healthy mice, where the proteasome presumably functions efficiently^14, 15, 35^. Yet, an open question is whether this same system is equally effective at regulating protein abundance in aged mice and mice undergoing active retinal degeneration, which is arguably a more relevant physiologic context for this approach. Therefore, in this study, we evaluated how well the DHFR DD system functioned in the retinas of aged (up to 24 mo) Balb/c and C57BL6/J mice, and in environmental (light-induced retinal degeneration) and genetic (*rd2*, *Abca4^−/−^*) models of retinal degeneration. We found that the DHFR DD system can be used effectively in all ages of mice and models tested, establishing a strong foundation for using the DHFR DD in eyes of a wide spectrum of healthy and diseased rodent models.

## RESULTS

### The DHFR DD is turned-over efficiently and stabilized effectively in the retina of aged mice

Age is the primary risk factor for the development of age-related macular degeneration (AMD), the leading cause of blindness in the elderly populace in industrialized nations^36^. Accordingly, certain aged animal models can be utilized to study the potential underpinnings of AMD^37^ and other age-associated diseases^37, 38^. Aging is typically associated with a decrease in protein quality control^39^ and a concomitant increase in cellular dysfunction/stress^40^. Whereas previous studies have indicated that proteasome activity decreases with age in a number of tissues^25–27^, human AMD donor eyes demonstrated increased chymotrypsin-specific proteasome activity in the retina with disease progression^41^. Thus, it is still unclear whether proteasome activity changes in the retina of aged vs. young subjects. Accordingly, we assessed proteasomal activity and the DHFR DD system in the retina of young (2 mo), adult (12 mo), and aged (17-24 mo) Balb/c mice (Fig. 1). Using fluorescent 7-amino-4-methylcoumarin (AMC)-conjugated polypeptides as substrates^42^, we measured the three proteolytic activities of the proteasome (i.e., chymotrypsin, trypsin and caspase). Surprisingly, all three proteolytic activities were unchanged in posterior eyecups of mice, regardless of age (Fig. 1A).

**Fig. 1.**
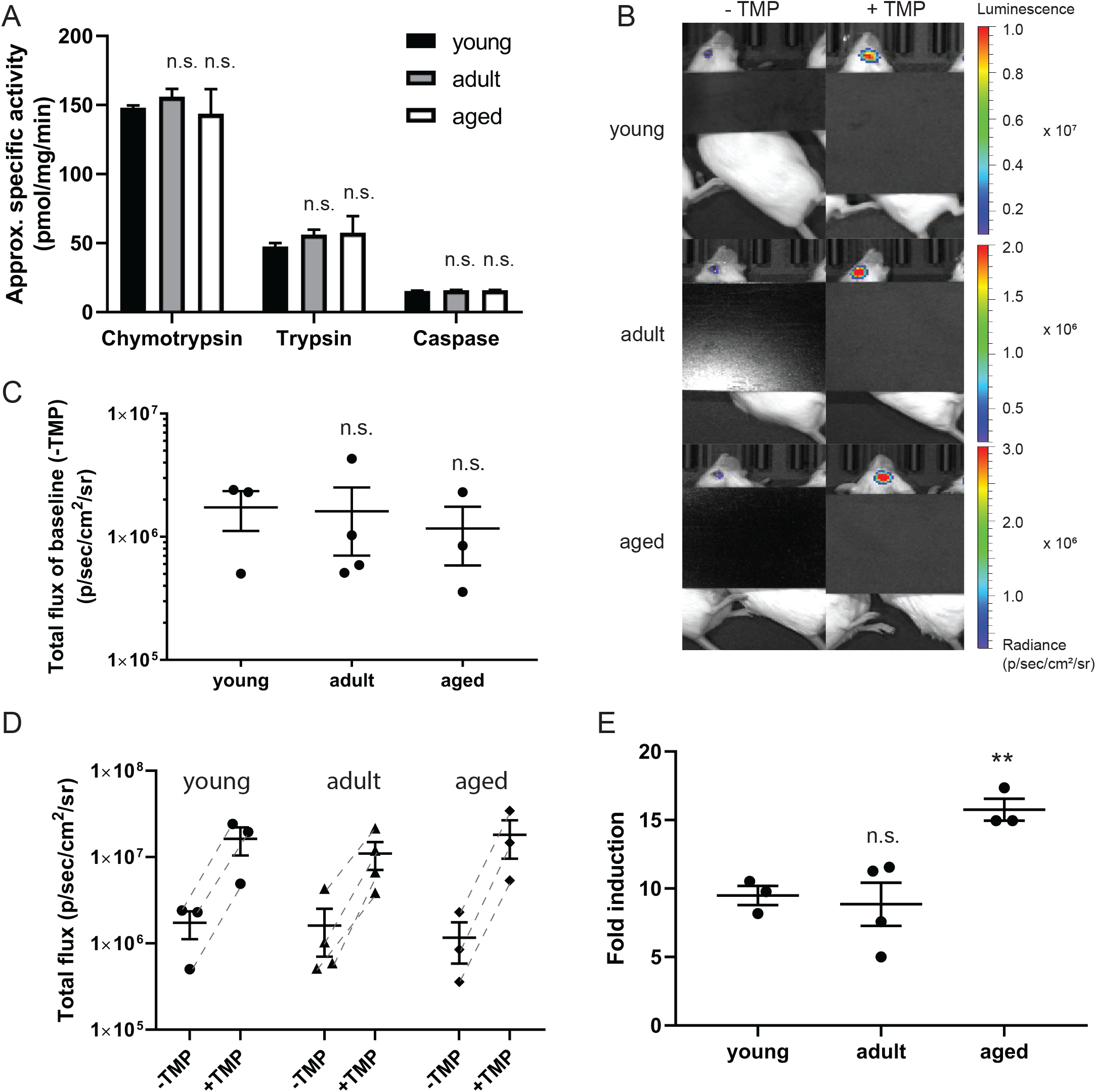
The DHFR DD is turned over efficiently and stabilized effectively in the retina of aged Balb/c mice. (A) Bar chart showing the three specific proteolytic activities (chymotrypsin, trypsin, and caspase activity) in the posterior eyecup of Balb/c mice at 2 mo (young), 12 mo (adult), and 17-24 mo (aged). Data are presented as mean ± standard deviation (SD) of n=3. (B) Representative images of bioluminescence signals originating from intravitreally injected mice at the indicated ages before (“− TMP”) and 6 h after TMP treatment (1 mg/mouse via gavage, “+ TMP”. n = 3 mice per age group). (C) Quantitation of the basal bioluminescence signal in the eye before TMP treatment. (D) Total flux numbers of the mice plotted before and after TMP treatment. (E) Fold induction of the bioluminescent signal of each mouse is plotted. For panels (C-E), data are represented as mean ± SEM. Statistical analysis in panels (A), (C) and (E) was conducted by unpaired, 2-tailed *t*-test assuming equal variance compared to 2 mo samples, n.s., not significant, ** p < 0.01. Note: a black tarp is used in (B) to block apparent background luminescence that can originate from the ear tag of the mice.

Next, we evaluated if the DHFR DD could function as effectively in aged mouse retinas compared to young mouse retinas in promoting protein degradation in the absence of the stabilizer, TMP. rAAV2/2[MAX], a serotype which transduces the entire neural retina after intravitreal injection^43, 44^, encoding DHFR.firefly luciferase (DHFR.FLuc), was intravitreally injected into young, adult and aged Balb/c mice. Based on the observed proteasome activities (Fig. 1A), we hypothesized that DHFR.FLuc should be turned-over efficiently across all ages, yielding a similar basal bioluminescent signal in the absence of TMP. Indeed, similar baseline signals were observed in all age groups prior to TMP administration (“− TMP”, Fig. 1B, C). Moreover, TMP effectively stabilized DHFR.FLuc in all ages of mice (“+ TMP”) (Fig. 1B, D-E). Interestingly, the aged mice demonstrated a significantly higher fold induction (15.8 ± 0.6 fold) in bioluminescence after TMP administration when compared to young mice (9.5 ± 0.5 fold, Fig. 1E, p ≤ 0.01, unpaired, 2-tailed *t*-test).

We paralleled our Balb/c aged mouse experiments in a separate genetic background, the C57BL6/J pigmented mouse. In an assay designed to complement the proteasome activity assay (e.g., Fig. 1A), we initially assessed levels of polyubiquitin as well as two substrates, rhodopsin (ubiquitin-dependent degradation^45^), and p53 (ubiquitin-independent degradation^46^) in adult (8-12 mo) and aged mice (18-22 mo, Fig. 2A). We found no demonstrable difference in any of these client proteins, suggesting intact and functional degradation capabilities in these mice, reinforcing our observations in aged Balb/c mice. Next, adult and aged mice were intravitreally injected as described above and tested for baseline and induced DHFR.FLuc bioluminescence. In this pigmented background, the total flux (i.e., bioluminescence) is ~2 orders of magnitude lower than that observed in non-pigmented mice due to absorbance of the luminescence by pigment. Nonetheless, as observed with Balb/c mice, similar baseline signals were recorded in adult compared to aged mice (Fig. 2B, C). TMP administration promoted DHFR.FLuc stabilization in both ages of mice (Fig. 2D), although as we observed with Balb/c mice (Fig. 1E), a significant increase in TMP-mediated fold induction was measured in aged mice (9.3 ± 0.6 fold) vs. adult mice (5.9 ± 0.9 fold, Fig. 2E, p ≤ 0.01, unpaired, 2-tailed *t*-test). While largely speculative, possible explanations for this observed increase could be due to age-related differences in TMP accessibility to the retina due to blood retinal barrier breakdown with age^47^ or altered TMP pharmacokinetics in older mice.

**Fig. 2.**
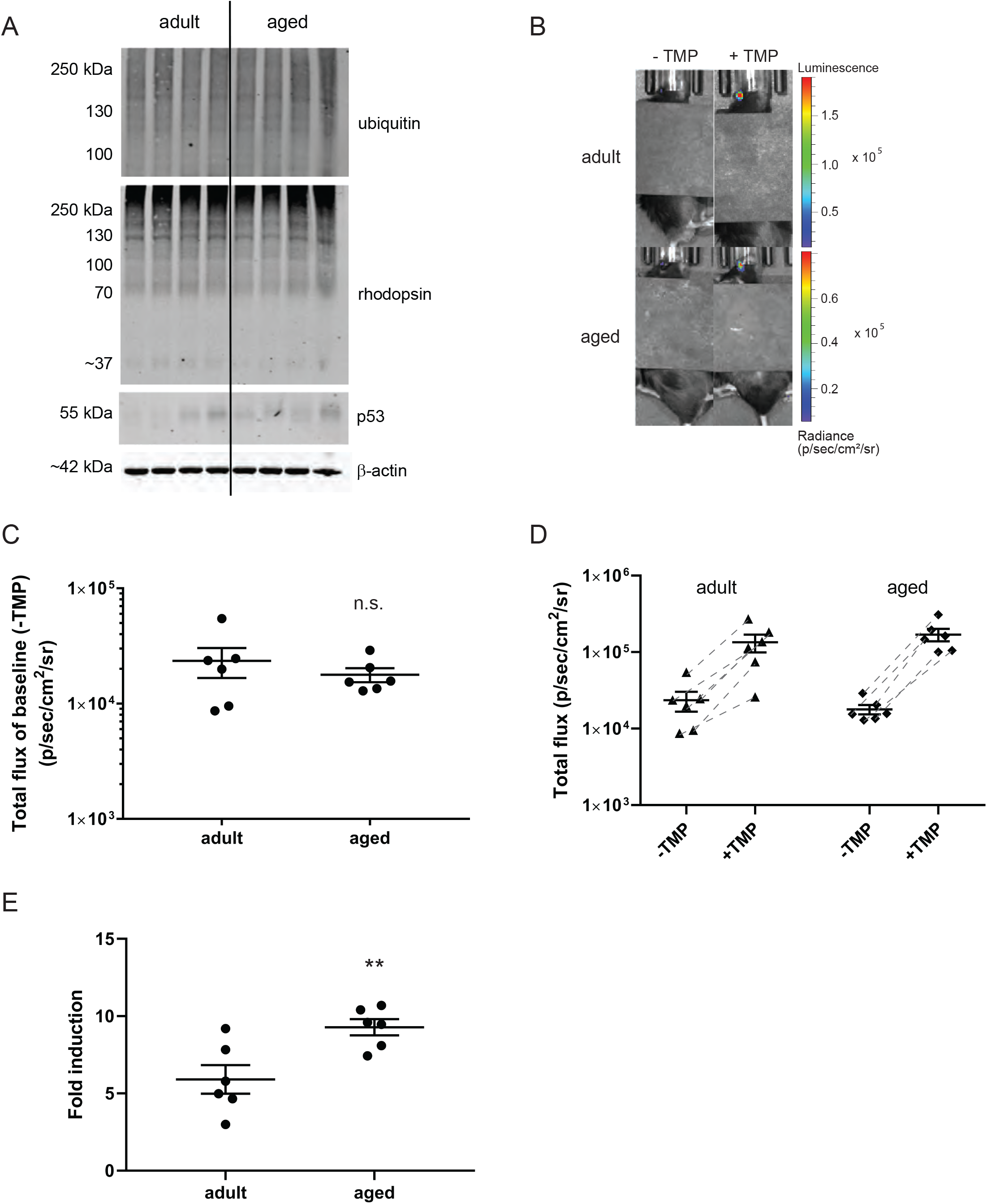
The DHFR DD is degraded efficiently and stabilized readily in the retina of aged C57BL6/J mice. (A) Western blot of adult (12 mo) and aged (20 mo) C57BL6/J mice posterior eyecups using ubiquitin, rhodopsin, p53 and β-actin antibodies (n=4). (B) Representative bioluminescent images of intravitreally-injected, untreated (“− TMP”) and treated (“+ TMP”, 1 mg/mouse, imaged 6 h post gavage, n = 6 mice per age group). (C) Quantitation of the basal bioluminescence signal in the eye before TMP treatment. (D) Total flux numbers of the mice plotted before and after TMP treatment. (E) Fold induction of the bioluminescent signal of each mouse is plotted. For panels (C-E), data are represented as mean ± SEM. Statistical analysis in panels (C) and (E) was conducted by unpaired, 2-tailed *t*-test assuming equal variance, n.s., not significant, ** p < 0.01. Note: a black tarp is used in (B) to block apparent background luminescence that can originate from the ear tag of the mice.

These DD-based bioluminescence findings in adult and aged mice were additionally verified in separate experiments employing different DHFR DD, DHFR fused to yellow fluorescent protein (DHFR.YFP, Sup. Fig. 1A-D). Moreover, we verified that the observed increase in bioluminescent FLuc signal did not originate from changes in FLuc mRNA transcript levels after TMP stabilization, but rather from stabilization at the protein level (Sup. Fig. 2). Overall, the combination of these bioluminescent and western blotting data demonstrate that multiple DHFR DDs can be efficiently degraded in the absence of TMP while stabilizable in the presence of TMP, regardless of mouse age.

### The DHFR DD is degraded and stabilized effectively in the retina of a light-induced retinal degeneration mouse model

Environmental factors, including smoking, diet, and possibly sunlight exposure, have been shown to contribute to the risk and progression of retinal diseases such as AMD, likely by increasing oxidative stress^48, 49^. Light-induced retinal degeneration (LIRD) is a rapid and aggressive method to induce retinal degeneration^50^ partly through the generation of oxidative stress^51^, photoreceptor apoptosis^52^ and/or inflammation in the retina^53^. We induced retinal degeneration using the LIRD method described previously^54^ by intraperitoneal injection of fluorescein and subsequent exposure to 50,000 lux of light for 3 min. Fundus images were obtained before (“− Light”) and 2 days after model generation (“+ Light”), showing severe pigmentary changes in a circular area that corresponds to the light exposure area around the optic nerve head post damage (Fig. 3A), indicating retinal degeneration (previously verified in more detail)^54^. We next measured the proteasome activity in LIRD mice 1 week after light injury and in age-matched, young untreated control mice. Although enhanced apoptosis and the expression of oxidative stress response genes are involved in the LIRD mouse model^54^, individual proteasome enzyme activities were not significantly different between LIRD mice and control mice (Fig. 3B). Furthermore, probing of select substrates (polyubiquitination, rhodopsin, p53) using western blotting corroborated the proteasome activity findings, demonstrating no appreciable differences between control and LIRD mice (Fig. 3C).

**Fig. 3.**
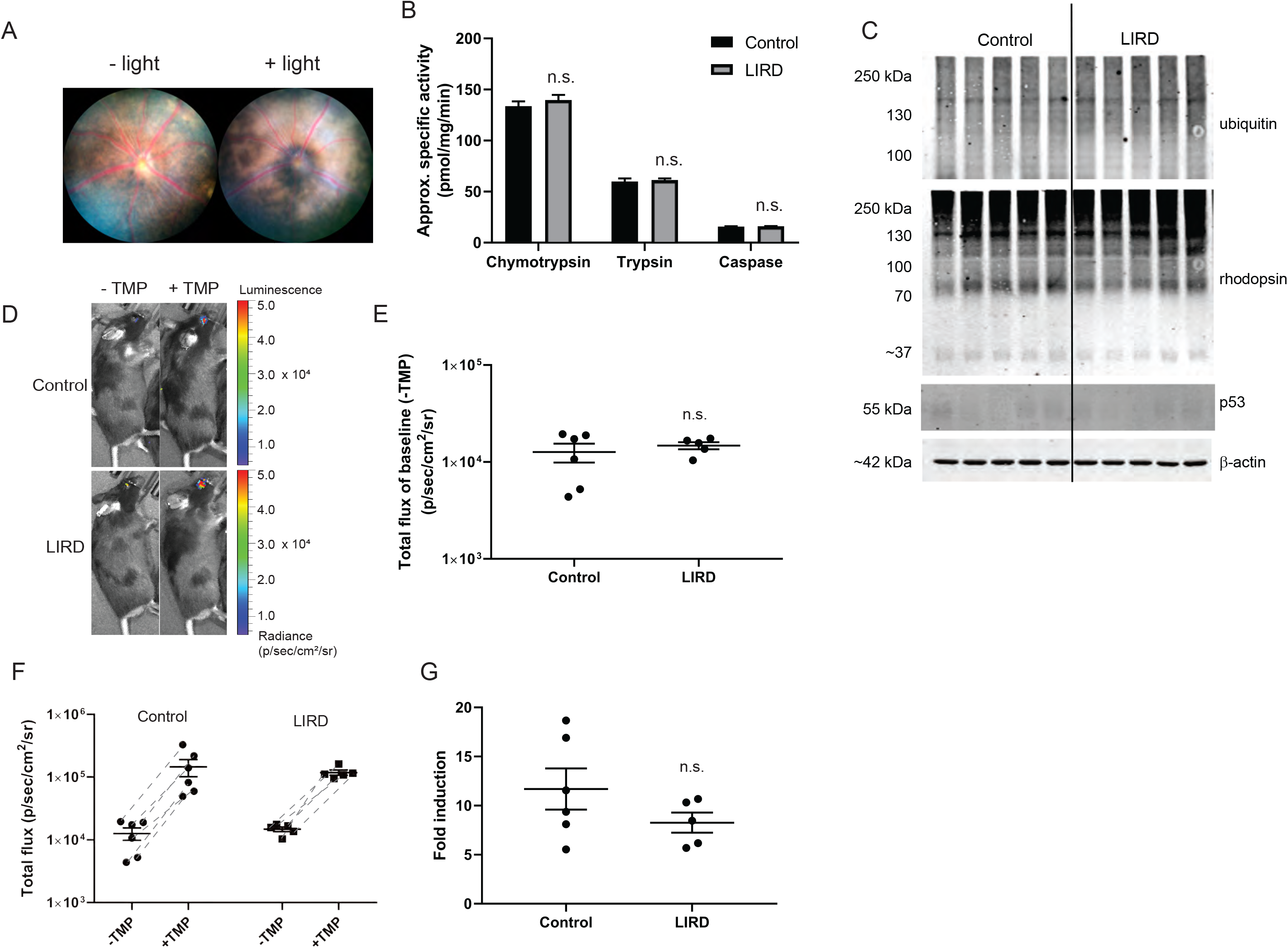
Light-induced retinal degeneration (LIRD) mice maintain the ability to degrade and stabilize the DHFR DD. (A) A representative fundus image before (“− Light”) and after (“+ Light”) LIRD (n ≥ 5). (B) Proteasome enzyme activities in the posterior eyecups of age-matched WT C57BL6/J mice (“control”) and LIRD mice harvested 1 week post light injury. Data are represented as mean ± SD, n=3. (C) Western blot of control and LIRD mouse posterior eyecups using ubiquitin, rhodopsin, p53 and β-actin antibodies (n=5). (D) Bioluminescence imaging of representative control and LIRD mice before (“− TMP”) and 6 h after TMP treatment (1 mg/mouse via gavage, “+ TMP”, n ≥ 5). (E) Quantitation of basal bioluminescence signal before TMP treatment in control and LIRD mice. (F) Total flux numbers of control mice and LIRD mice before and after TMP treatment. (G) The fold induction of the signal of each mouse. Data are represented as mean ± SEM in panels (E-G). Statistical analysis in panels (B), (E), (F), and (G) was conducted by unpaired, 2-tailed *t*-test assuming equal variance compared to control mice. n.s., not significant.

Given these promising results, we next evaluated whether mice undergoing LIRD were also able to effectively control DHFR DD abundance as well as control mice. We intravitreally injected C57BL6/J mice AAV encoding for DHFR.FLuc and facilitated AAV expression in the eye for 10 days prior to LIRD as described above. Both sets of mice demonstrated a similar ability to degrade the DHFR DD (Fig. 3D, E). Moreover, there were no significant differences in TMP-induced bioluminescence signals (Fig. 3D, F, G); control mice showed an 11.7 ± 2.1 fold induction whereas LIRD mice displayed a 8.3 ± 1.0 fold (Fig. 2G), revealing that the DHFR DD was degraded and stabilized effectively in mice actively undergoing LIRD.

### The DHFR DD is readily degraded and effectively stabilized in the retina of genetic retinal degeneration mouse models

A number of naturally-occurring and genetically-engineered mouse strains serve as surrogates of human diseases like retinitis pigmentosa (*e.g*., the *rd2* mouse, a surrogate for human peripherin mutations that cause retinitis pigmentosa^55–57^) and Stargardt disease (*e.g*., the *Abca4^−/−^* mouse as a Stargardt disease model^58^). Consequently, these models have been useful in studying disease mechanisms and therapy development. Accordingly, we tested the regulation of the DHFR DD system in these genetic retinal degeneration mouse models in the hope of eventually applying this conditional gene regulation approach for the treatment of disease.

*Rd2* mice lack the Rds/peripherin gene, which encodes a glycoprotein that is required for the formation of photoreceptor outer segment discs^55, 59, 60^. Homozygous *rd2* mice show an early onset of photoreceptor degeneration, which we confirmed by histology indicating the loss of outer segment formation by 1 mo of age (Fig. 4A). Homozygous *rd2* mice demonstrated significantly reduced levels of chymotrypsin (~40%, Fig. 4B, p ≤ 0.05, unpaired, 2-tailed *t*-test), the predominant enzyme activity of the proteasome. Moreover, when we examined the steady state levels of particular protein substrates by western, we detected a virtual absence of rhodopsin, presumably due to photoreceptor degeneration, but hinting at a robust level of degradative capacity still present (Fig. 4C). To examine if the reduced level of proteasome activity was still sufficient to degrade the DHFR DD, we compared the baseline bioluminescence signal generated from subretinal injection of DHFR.FLuc rAAV2/2[MAX] before TMP stabilization. Surprisingly, we found that the basal signal, which implies how readily the DHFR DD is turned over, was significantly lower in *rd2* mice than controls (Fig. 4D, E), which is the opposite of what we would have predicted based on the chymotrypsin findings (Fig. 4B). This is possibly due to loss of photoreceptors that are expressing the DHFR DD during degeneration. Nonetheless, both WT agouti and *rd2* mice produced more bioluminescent signal after TMP treatment (Fig. 4D, F); the fold induction of bioluminescent signal was lower in the *rd2* mice (7.1 ± 0.8 fold [*rd2*] vs. 10.8 ± 1.5 fold [WT]), but did not reach statistical significance (Fig. 4G), suggesting the TMP-induced stabilization of the DHFR DD system in the two mouse groups is similar.

**Fig. 4.**
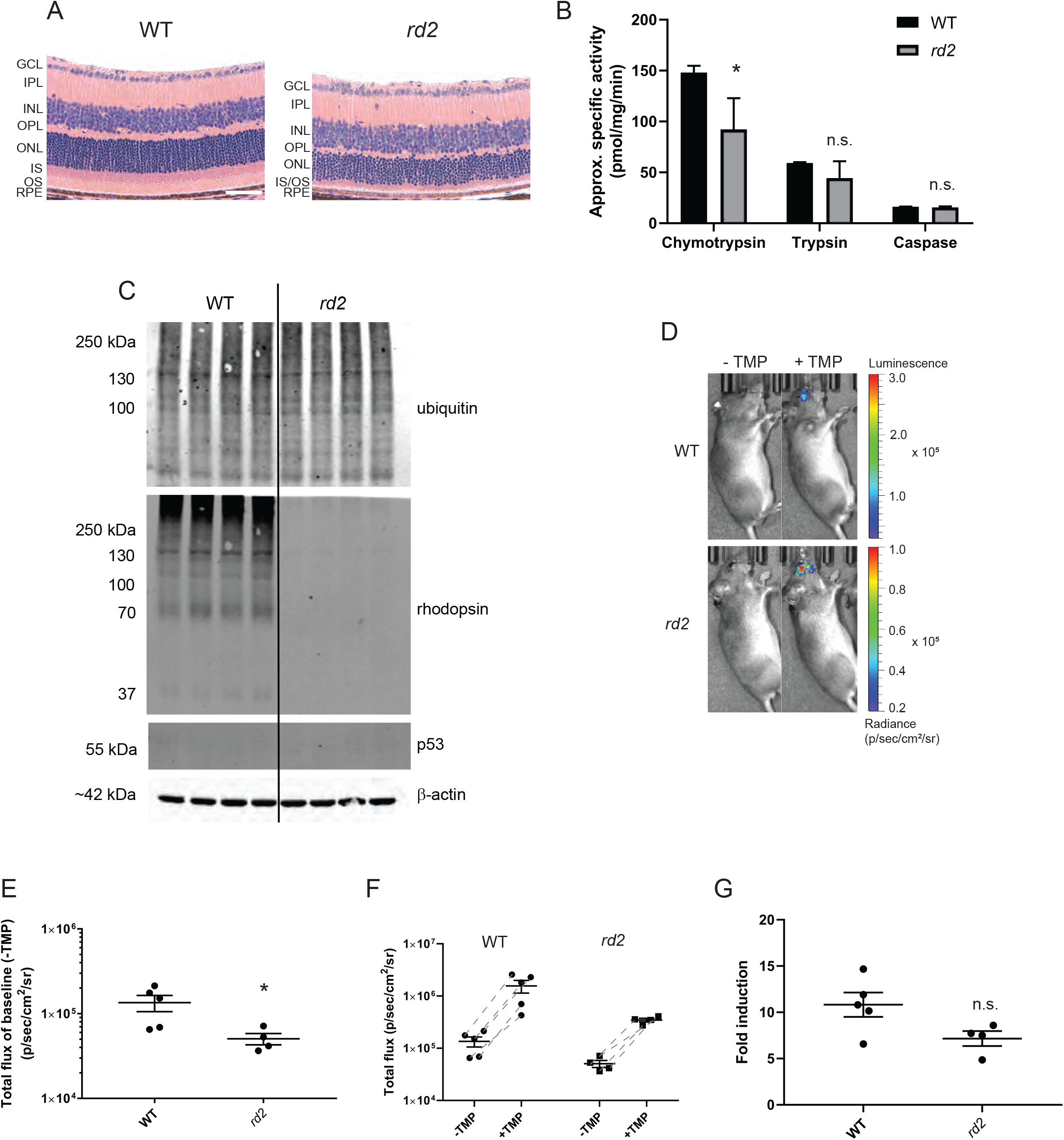
The DHFR DD is readily degraded and effectively stabilized in retinitis pigmentosa model mice. (A) H&E staining showing the retinal cell layers of 1 mo WT agouti and homozygous *rd2* mice. RPE: retinal pigment epithelium, OS: outer segment; IS: inner segment; ONL: outer nuclear layer; OPL: outer plexiform layer; INL: inner nuclear layer; IPL: inner plexiform layer, GCL: ganglion cell layer. Scale bar = 50 μm. (B) Posterior eyecup proteasome enzyme activities of 3 mo WT and *rd2* mice, presented as mean ± SD of n=3. (C) Western blot of 1 mo WT and *rd2* mouse posterior eyecups using ubiquitin, rhodopsin, p53 and β-actin antibodies (n=4). (D) Representative bioluminescence images from subretinally-injected mice at 1 mo before (“− TMP”) and after (“+ TMP”) TMP treatment, n ≥ 4. (E) The total flux of basal signal before TMP treatment. Total flux of the ocular bioluminescent signal was quantified and plotted. (G) Quantitation of bioluminescence imaging represented as fold induction. In panels (E-G), data are represented as mean ± SEM. Statistics in panels (B), (E), and (G) is analyzed by unpaired, 2-tailed *t*-test assuming equal variance, * p < 0.05, n.s., not significant.

In addition to the *rd2* mice, we also evaluated the induction of DHFR.FLuc in the *Abca4^−/−^* mouse, a model of autosomal recessive Stargardt disease^61–63^. These mice lack the Abca4 phospholipid ATPase transporter (located in the photoreceptors and RPE) and are characterized by a progressive increase in bisretinoid autofluorescence and accompanying oxidative, inflammatory, and complement-related stress^64–66^, but do not typically show overt histological disruptions of the retina, even by 6 mo of age (Fig. 5A). As we have performed with the three previous model systems, proteasome activities and select protein clients were measured in WT Balb/c and *Abca4^−/−^* mice as described above (Fig. 5B, C), with no appreciable differences aside from sightly elevated levels of p53 in WT Balb/c mice. In a separate cohort of rAAV2/2[MAX]-intravitreally-injected mice (injected at ~2 mo, and allowed to age until 6 mo prior to imaging), the basal bioluminescence signal was quantified in *Abca4^−/−^* mice and age-matched WT Balb/c mice (Fig. 5D, E). While there is a significant increase in the A500 nm A2E precursor and A2E in *Abca4^−/−^* mice at this time point^66^, which is corroborated by autofluorescent retinal pigmented epithelium (RPE) flat mounts (Sup. Fig 3), the extent of retinal degeneration is not substantial at this age^67^ (Fig. 5A). Therefore, it was not particularly surprising to observe similar basal bioluminescent signals (Fig. 5E) and that TMP could stabilize DHFR.FLuc well in both mouse lines. However, when quantified as fold change in bioluminescent signal, the *Abca4*^−/−^ mice actually showed a significantly decreased induction (13.1 ± 0.9 fold [WT] vs. 9.5 ± 0.8 fold [Abca4^−/−^], Fig. 5G, p ≤ 0.05, unpaired, 2-tailed *t*-test). Such a minor difference (~27%) would not prevent the application of the DHFR DD system to this mouse model, but should be a consideration going forward. Nonetheless, the culmination of these observations validate and extend the use of the DHFR DD system in a range of distinct mouse models of retinal degeneration.

**Fig. 5.**
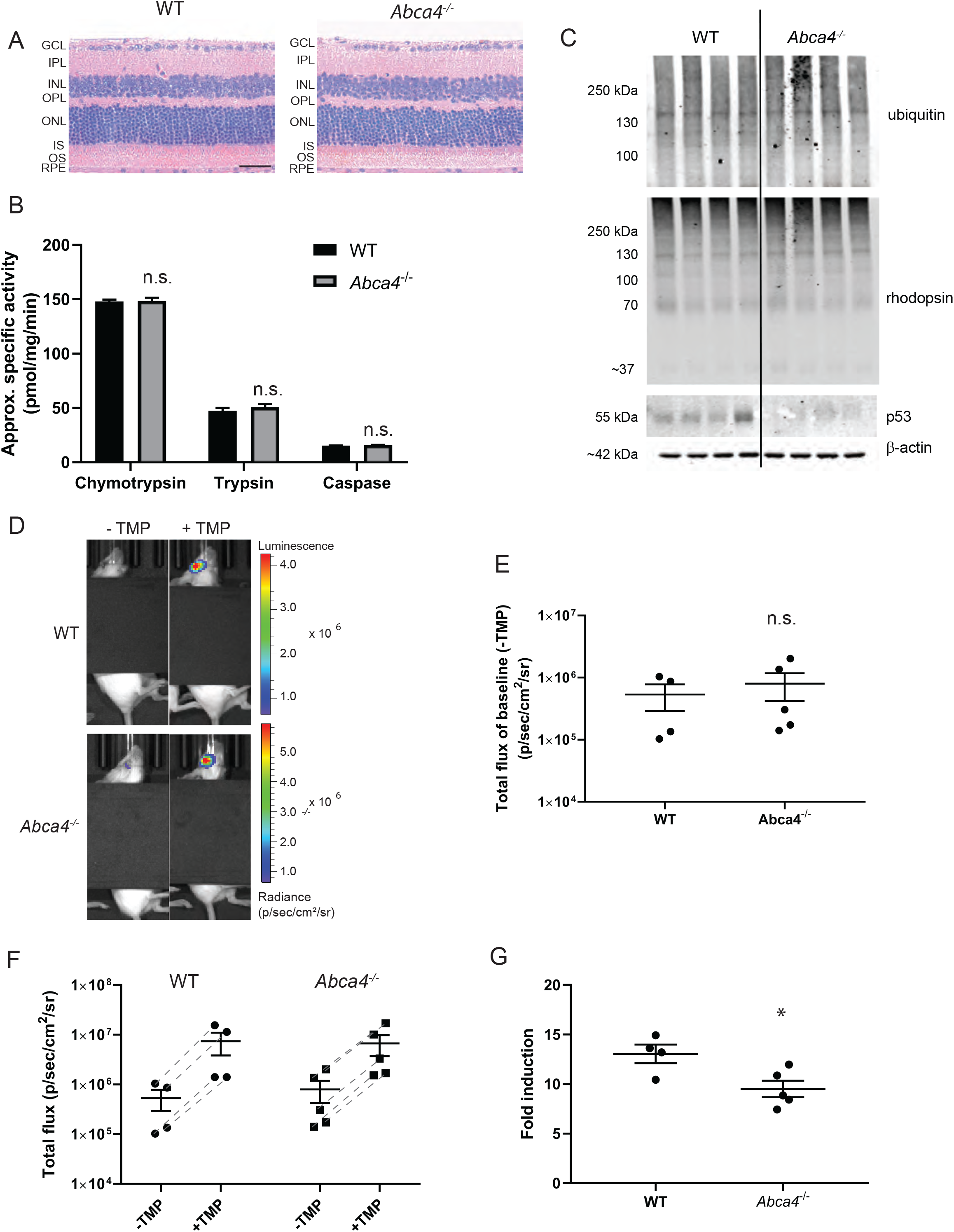
The DHFR DD can be effectively used in a mouse model of Stargardt disease. (A) H&E staining showing the retinal cell layers of 2 mo WT and *Abca4*^−/−^ mice. RPE: retinal pigment epithelium, OS: outer segment; IS: inner segment; ONL: outer nuclear layer; OPL: outer plexiform layer; INL: inner nuclear layer; IPL: inner plexiform layer, GCL: ganglion cell layer. Scale bar = 50 μm. (B) The three specific proteasome activities were measured in the posterior eyecups of aged-matched WT and *Abca4*^−/−^ 2 mo mice. Data are represented as mean ± SD of n=3. Note: WT mice in this panel are the same as the “young” mice in Fig. 1A. (C) Western blot of WT and Abca4^−/−^ 6 mo posterior eyecups using ubiquitin, rhodopsin, p53 and β-actin antibodies, n=4. (D) Representative ocular bioluminescence signals before and after TMP treatment. (E-G) Quantitation of bioluminescence imaging plotted as basal signal (E), total flux values of control or *Abca4*^−/−^ mice plotted before (“− TMP”) and after TMP (“+ TMP”, F), or fold induction of bioluminescent signal of each mouse (G). Data in panels (E-G) are presented as mean ± SEM. Statistical analysis was conducted by unpaired, 2-tailed *t*-test assuming equal variance compared to WT mice, * p < 0.05, n.s., not significant. Note: a black tarp is used in (D) to block apparent background luminescence that can originate from the ear tag of the mice.

## DISCUSSION

Until now, a thorough characterization of the regulation of the DHFR DD strategy for regulating protein abundance in the retina was only tested in young, healthy mice^44^. In this study, we extended these observations to non-pigmented and pigmented aged mice as well as a series of environmental/genetic retinal degeneration mouse models. We found that for the most part, it appears that proteasome activity is largely maintained across these model systems, but that in certain forms of aggressive retinal degeneration, such as the homozygous *rd2* mouse, proteasome activity may be compromised. Similar findings of proteasome overload in various forms of retinal degeneration have also been observed (*rd2* [a.k.a., Rds], and retinitis pigmentosa caused by the P32H mutation in rhodopsin)^34, 68^. Nonetheless, even with lower proteasome enzyme activity levels observed in *rd2* mice, we never observed higher basal levels of DHFR.FLuc bioluminescence under any conditions, suggesting that there is a fair degree of leeway in retinal proteolytic capacity that affords efficient degradation of the DHFR DD. One important limitation of this study is that we were only able to explore a handful of representative retinal degeneration mouse models, which do not cover the entire pathological spectrum of potentially confounding factors. Moreover, except for the aged mice, we evaluated mice at a single time point during the retinal degeneration, which fails to account for additional physiological changes that may occur as disease progresses. Future longitudinal studies could shine light onto the long-term stability and biological tolerance for the DHFR DD.

Even with these limitations, the DHFR DD system appears to be increasingly flexible with respect to diversity of model systems to which it can be applied (e.g., yeast^69^, mice, drosophila^70^, C. elegans^71^), the proteins it can effectively regulate^11, 72–75^, and at the level of the small pharmacologic chaperone^13–15, 76^. With this increasing flexibility comes the ability to apply and fine-tune the DHFR DD system to achieve higher degrees of spatial (guided by AAV or virus tropism and/or a cell-type specific promoter) and temporal (timing of small molecule administration and/or pharmacokinetic properties) control over protein abundance, and therefore cellular signaling; an exciting prospect for customizable or personalized gene therapy approaches.

In summary, our work demonstrates the efficacy of the DHFR-based system for conditional regulation of protein abundance in aged mice and representative retinal degeneration mouse models, laying the foundation for the application of the DHFR DD system as a biological probe or gene therapy strategy for a variety of distinct mouse models of eye disease.

## METHODS

### Mouse use

All animal experiments were performed according to the guidelines of the Association for Research in Vision and Ophthalmology (ARVO) Statement for the Use of Animals in Ophthalmic and Vision Research and were approved by the Institutional Animal Care and Use Committee (IACUC) of UT Southwestern Medical Center, Dallas, TX, USA. WT Balb/c mice originated from heterozygous breeding schemes from R345W^+/−^ EFEMP1 mice (courtesy of Dr. Lihua Marmorstein, private stock at The Jackson Laboratory, Bar Harbor, ME). WT C57BL6/J control and experimental mice used for light-induced retinal degeneration experiments were purchased from the UT Southwestern Mouse Breeding Core and were genotyped to confirm the absence of the potentially confounding *rd8* mutation^77^. WT adult (8-12 mo), aged (20-22 mo) C57BL6/J mice, *rd2* (stock # 001979^78^), and WT agouti (stock # 001912) mice were also obtained from The Jackson Laboratory. *Abca4^−/−^* mice in the Balb/c background^64^ were obtained from Dr. Roxana Radu, Jules Stein Eye Institute, University of California Los Angles. Equal numbers of age-matched, littermate male and female mice were used whenever possible. Mice were provided standard laboratory chow and allowed free access to water, in a climate-controlled room with a 12 h light/12 h dark cycle.

### Proteasome activity assay

Mice were euthanized by overdose of ketamine/xylazine (180 mg/kg, 24 mg/kg, respectively) and their eyes were enucleated to prepare posterior eyecup samples. The anterior segment of the eye, the optic nerve, and the muscles attached to the outside of the eyeball were discarded, and the remaining posterior eyecups from both eyes were snap-frozen in liquid nitrogen and stored at −80° until use. When ready to use, the posterior eyecup samples were homogenized on ice in 600 μL lysis buffer containing 50 mM Tris·HCl, pH 7.5, 40 mM KCl, 5 mM MgCl_2_, 1 mM DTT and 0.5 mM ATP. The crude cell lysate was centrifuged at 17,000 g for 20 min at 4° and the supernatant was collected. Protein concentration in the supernatant was quantified by a bicinchoninic acid (BCA) assay (Piece, Rockford, IL, USA). Note: the small amount of DTT in the lysis buffer does contribute to a minimal amount of BCA signal; therefore, the resulting specific activity of this assay is considered an approximation. Fifty μL of ~0.25 mg/mL protein was used to measure proteasome activity by using a proteasome activity assay kit (BioVision, Milpitas, CA, USA). All the components in the kit were used except for the provided substrate. Instead of using the provided substrate, separate substrates were purchased and used to evaluate each proteasomal protease activity. Final concentrations of 100 μM Suc-Leu-Leu-Val-Tyr-AMC (Cayman Chemical, Ann Arbor, MI, USA), Boc-Leu-Arg-Arg-AMC (AdipoGen, San Diego, CA, USA) or Ac-Nle-Pro-Nle-Asp-AMC (Cayman Chemical) were used as substrates to measure chymotrypsin-, trypsin- and caspase-specific proteolytic activity, respectively. A 1.5 h kinetics curve was monitored at 37° using a Synergy 2 multi-mode microplate reader (BioTek, Winooski, VT, USA). The absolute amount of cleaved substrate was calculated by an AMC standard curve. The assay was conducted with and without 100 μM MG-132 (Sigma-Aldrich, St. Louis, MO, USA) to confirm proteasomal specificity of the AMC generation. All proteasome assay data were performed in parallel across the model systems.

### Intravitreal Injections

Intravitreal injections were conducted on Balb/c, C57BL6/J, and Abca4^−/−^ mice following the protocol described previously^15, 35^. Mice were anesthetized with a ketamine/xylazine cocktail (120 mg/kg, 16 mg/kg, respectively) followed by pupillary dilation using cyclopentolate hydrochloride (1% [w/v]) (Alcon, Fort Worth, TX, USA) and tropicamide (1% [w/v]) (Alcon). GenTeal eye gel (severe dry eye formula, Alcon) was applied before injection to prevent corneal drying. A Stemi 305 stereo microscope (Zeiss, Oberkochen, Germany) was used to assist with the injection. The right eye was pierced by a 30G needle approximately 1 mm posterior to the supratemporal limbus. Then a 33G ½ needle fitted to a Hamilton micro-syringe (Hamilton, Reno, NV, USA) was inserted into the previous incision at a 45-degree angle until needlepoint was mid-vitreous. Two microliters of rAAV2/2[MAX]^43^ encoding for Nano luciferase 2A DHFR.FLuc (7 × 10^9^ viral genomes, prepared by the UNC Viral Vector Core, Chapel Hill, NC, USA) was slowly injected into the vitreous over the course of ~1 min. Note: Nano luciferase was not used in these studies because of issues with substrate administration and penetrance into the eye. Afterwards, the needle was held stable for an additional minute before slowly removed. The left eye was injected with a sham vehicle (HBSS with 0.14% Tween [HBSS-T]) using the same injection procedure. After injection, bacitracin zinc and polymyxin B sulfate ointment (Bausch & Lomb, Tampa, FL, USA) and GenTeal eye gel were applied to each eye. Mice were kept warm on a heating pad until regaining consciousness. For Balb/c mice shown in Fig. 1, and C57BL6/J mice shown in Fig. 2, mice at the indicated ages were injected with AAV ~10 days prior to initial baseline imaging and subsequent imaging post stabilization with TMP. Young LIRD mice were intravitreally injected with AAV 10 days prior to initiation of the LIRD protocol. *Abca4*^−/−^ mice were injected at ~2 mo with AAV and allowed to age out to 6 mo prior to baseline and stabilization readings.

### Subretinal injections

WT agouti and *rd2* mice at four days postnatal (P4) were anesthetized on ice until they stopped moving. Using a Stemi 305 stereo microscope to assist with the injection, the eyelid was slit by a 25G 5/8 needle to allow the eye to proptose. The right eye was pierced by a 30G needle approximately 1 mm posterior to the supratemporal limbus. Then a 33G ½ needle with blunt tip fitted to a Hamilton micro-syringe was inserted into the previous incision at a 45-degree angle until the needlepoint met resistance posterior to the retina. Half a microliter of rAAV2/2[MAX] encoding for DHFR.FLuc (1 × 10^9^ viral genomes) was injected into the subretinal space. The left eye was injected with a sham vehicle (HBSS with 0.14% Tween [HBSS-T]) using the same injection procedure. After injection, bacitracin zinc and polymyxin B sulfate ointment was applied to each eye. Mice were kept warm on a heating pad until regaining consciousness. Mice were imaged for baseline bioluminescence and stabilization at 1 mo of age.

### Bioluminescence imaging

To establish an initial baseline signal, mice were placed in an IVIS Spectrum In Vivo Imaging System (UT Southwestern Small Animal Imaging Resource, PerkinElmer, Waltham, MA, USA) under anesthesia by circulating isoflurane. Each mouse was injected intraperitoneally with 150 mg luciferin/kg body weight, and the eyes were imaged for bioluminescence over a 20-min time course with 1 min interval between every image. At times, a black tarp was used to cover the mice to prevent background signal due to an ear tag. The next day mice were given 1 mg TMP (~50 mg/kg depending on mouse weight). Specifically, TMP was dissolved in 20 μL DMSO and then diluted with 40 μL PEG400 (Fisher Scientific, Waltham, MA, USA), 4 μL Tween 80 (Fisher Scientific), 20 μL cremaphor (Sigma-Aldrich) and 116 μL of 5% dextrose (Fisher Scientific) and administered by oral gavage. Six hours after oral gavage, the mice were imaged again in the same manner as described for basal signal establishment. The total flux number at the peak of the kinetics and the image with the peak number were used for plotting and comparison between “− TMP” and “+ TMP”. While in this manuscript we have performed bioluminescence imaging on Balb/c, C57BL6/J and agouti mice, we should note that Balb/c and agouti mice are preferred due to absence or reduction in ocular pigmentation. C57BL6/J can be used, but typically yield much lower absolute basal bioluminescence.

### Western blotting

For protein analysis involving ubiquitin, rhodopsin, or p53, mouse posterior eyecups were lysed with radioimmunoprecipitation assay (RIPA) buffer (Santa Cruz, Dallas, TX, USA) supplemented with Halt protease inhibitor (Pierce, Rockford, IL, USA) and benzonase (Millipore Sigma, St. Louis, MO, USA) and homogenized on ice using a Bel-Art™ Micro-Tube Homogenizer (Fisher Scientific, Waltham, MA, USA, cat# 03-421-215). For experiments involving assessment of the DHFR.YFP protein, neural retina was peeled from the eye cup and homogenized as described above. After homogenization, samples were incubated on ice for 10 mins then spun at max speed (14,800 rpm) at 4°C for 10 min. The supernatant was collected and protein concentration was quantified via a bicinchoninic assay (BCA) assay (Pierce Thermo Scientific, Rockford, IL, USA). Approximately 30 μg of soluble supernatant was run on a 4-20% Tris-Gly SDS PAGE gel (Life Technologies) and transferred onto a nitrocellulose membrane using an iBlot2 device (Life Technologies). After probing for total protein transferred using Ponceau S (Sigma-Aldrich), membranes were blocked overnight in Odyssey Blocking Buffer (LICOR, Lincoln, NE, USA). Membranes were probed with rabbit anti-rhodopsin (1:1000; Cell Signaling Technology, Danvers, MA. USA cat# 14825S), mouse anti-p53 (1:1000; Cell Signaling Technology, cat# 2524S), mouse anti-ubiquitin (1:1000; Santa Cruz, Dallas, TX. USA, cat# sc-8017), mouse anti-HA (1:1500, Pierce, Waltham, MA, clone 2-2.2.14, cat# 26183), and mouse anti-β-actin (1:5000; Sigma- Aldrich, cat# A1978). All Western blot imaging was performed on an Odyssey CLx (LI-COR) and band quantification was performed using Image Studio software (LI-COR).

### Fundus camera-delivered light-induced retinal degeneration (FCD-LIRD) mouse model generation

Ten days after intravitreal AAV injection, a group of randomly picked mice was designated as control (virus injected, fluorescein-injected, no light-induced degeneration) while the rest were used for the severe FCD-LIRD model^54^, here after simply referred to as LIRD. An advantage of this specific method is that one can induce retinal damage in the common C57BL6/J background. As previously described^54^, LIRD-designated mice were dark adapted overnight, and the next morning anesthetized with a ketamine/xylazine cocktail (120 mg/kg, 16 mg/kg, respectively) followed by pupillary dilation using cyclopentolate hydrochloride (1% [w/v]) and tropicamide (1% [w/v]). GenTeal eye gel was also applied to prevent dry cornea. After a clear fundus image was taken, mice were injected with fluorescein (2 mg, i.p.), incubated for 10 mins, and subsequently exposed to 50,000 lux of light for 3 min (continuously). Afterwards, bacitracin zinc and polymyxin B sulfate ointment was applied to the eye, and mice were kept warm on a heating pad until regaining consciousness. Two days after model generation, the fundus images of the LIRD-designated eyes were obtained to confirm retinal degeneration.

### Hematoxylin and Eosin (H&E) Staining

WT agouti mice and *rd2* mice at 1 mo were euthanized by ketamine/xylazine overdose. WT Balb/c and *Abca4^−/−^* mice at 6 mo were euthanized similarly. Eyes were rapidly enucleated and processed for freeze substitution following previous literature recommendations^79^ with slight modification. Enucleated eyes were submerged in chilled isopentane for 1 min, transferred to chilled 97% methanol and 3% acetic acid (MAA) and stored at −80° for 48 h. Then the vial was warmed stepwise to −20° for 24 h, followed by 4° for 4 h, and then room temperature for 2 h. MAA was then replaced with 100% ethanol and the eye was processed into paraffin-embedded sections. H&E staining was performed by the UT Southwestern Histo Pathology core and images of the sections were taken at 40× magnification using an inverted epifluorescent microscope (Zeiss).

## ACKNOWLEDGEMENTS

JDH is supported by an endowment from the Roger and Dorothy Hirl Research Fund, a vision research grant from the Karl Kirchgessner Foundation, a Macular Degeneration Research Grant from the BrightFocus Foundation (M2018099), a National Eye Institute R21 Grant (EY028261), a National Eye Institute R01 Grant (EY027785) and a Career Development Award from Research to Prevent Blindness. Additional support was provided by a National Eye Institute Visual Science Core Grant (P30 EY030413) and an unrestricted grant from RPB (both to the UT Southwestern Department of Ophthalmology).

## SUPPLEMENTAL FIGURE LEGENDS

**Sup. Fig. 1.**
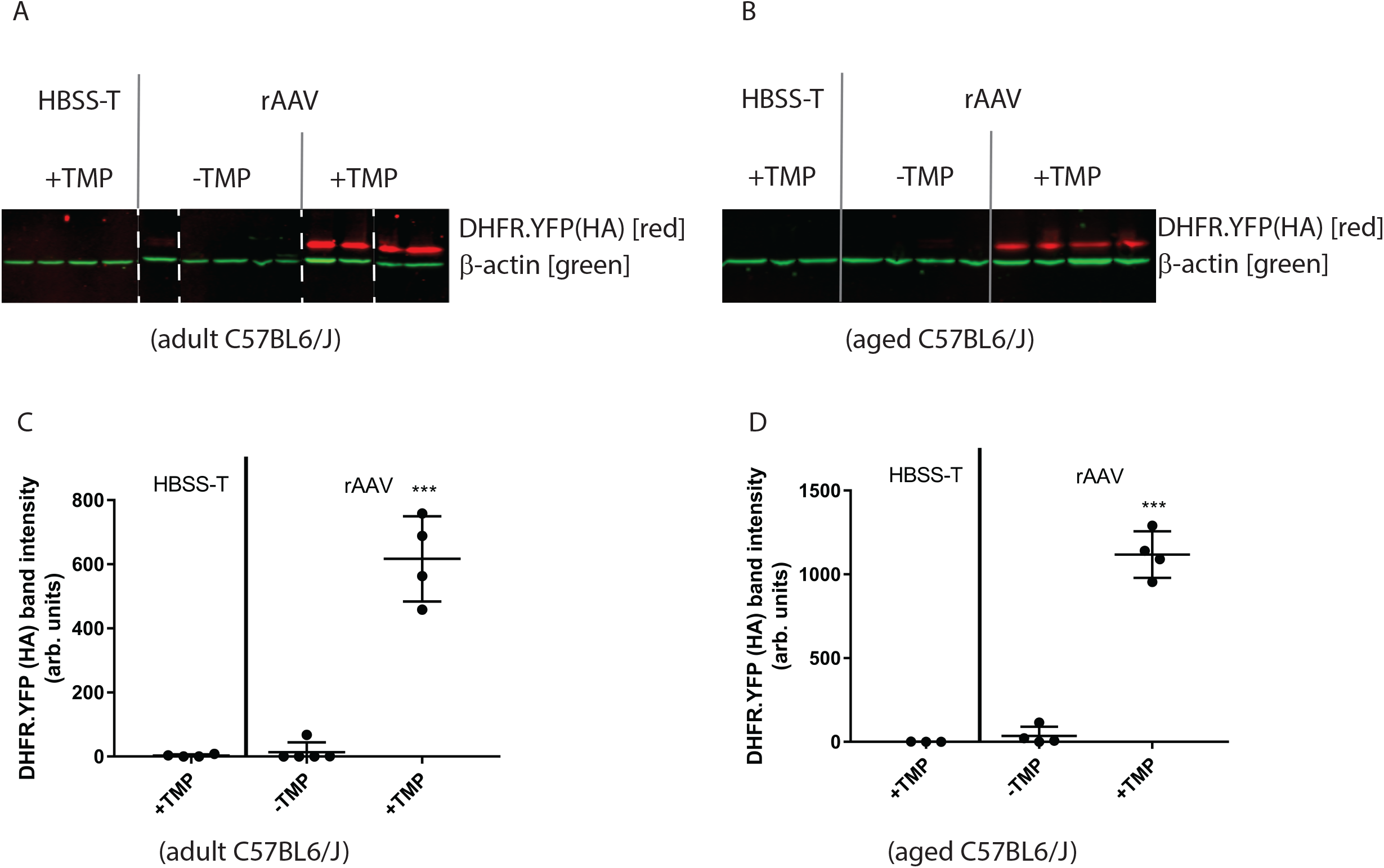
DHFR.YFP protein abundance is well regulated in the retina of adult and aged C57BL6/J mice. (A, B) adult (9 mo) or aged (18 mo) C57BL6/J mice were intravitreally injected with either Hanks buffered salt solution with 0.14% v/v Tween-20 (HBSS-T) or rAAV2/2[MAX] encoding for DHFR.YFP tagged C-terminally with a hemagglutinin (HA) and given normal water (“− TMP”) or water supplemented with TMP (“+ TMP”, 0.4 mg/mL, a saturated amount) in their drinking water overnight. Neural retina were isolated and analyzed by western blotting using HA and β-actin antibodies (n ≥ 4). (C, D) Quantification of HA band intensities shown in A, B. Statistical analysis was conducted by unpaired, 2-tailed *t*-test assuming equal variance compared to “− TMP” rAAV samples n.s., not significant, *** p < 0.001.

**Sup. Fig. 2.**
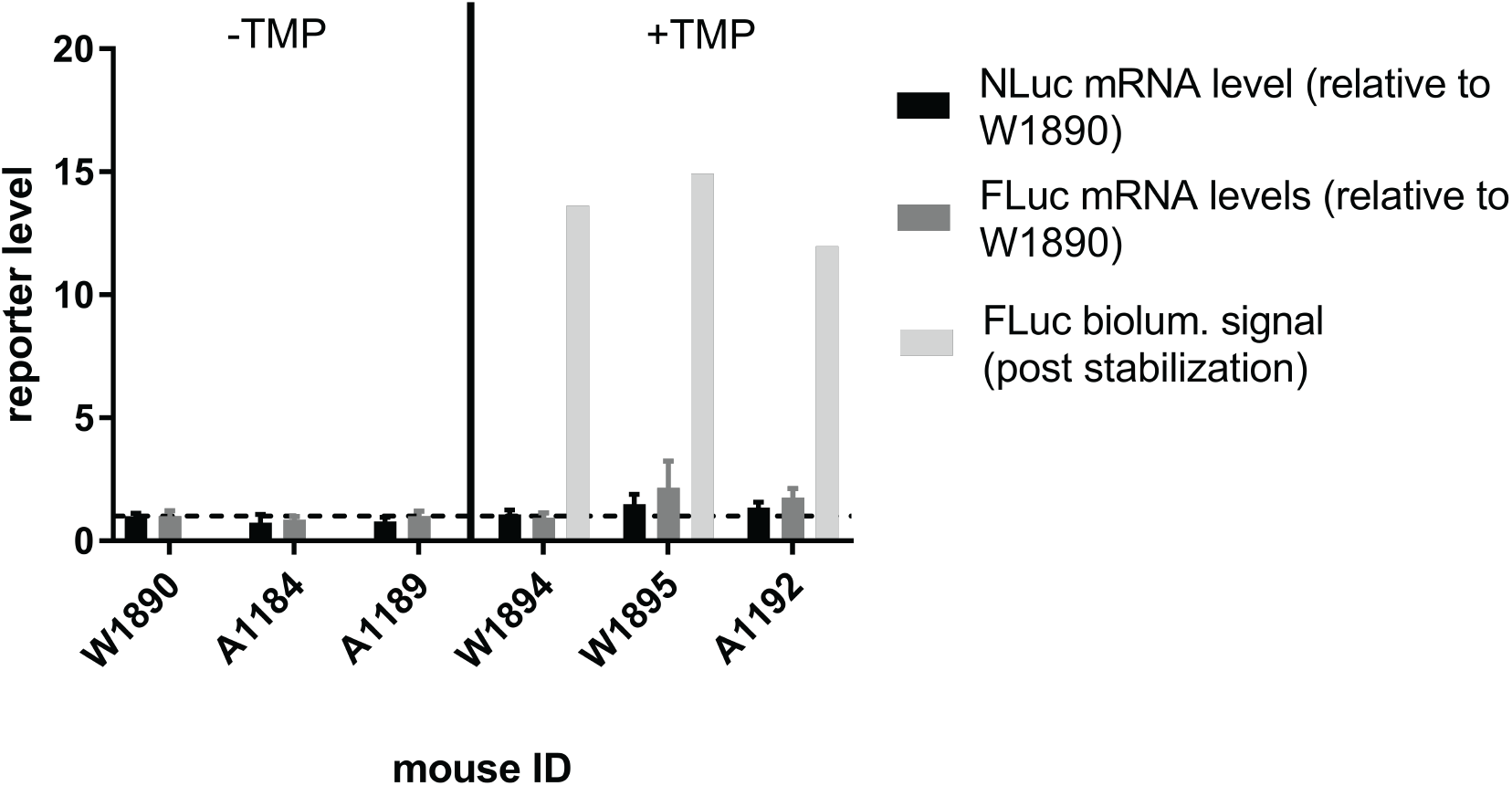
DHFR.FLuc is stabilized at the protein, not mRNA level. A series of young WT Balb/c (mouse IDs: W1890, W1894, W1895) or Abca4^−/−^ mice (mouse IDs: A1184, A1189, A1192) were intravitreally injected rAAV2/2[MAX] encoding for NLuc 2A DHFR.FLuc. A portion of the mice remained untreated (“− TMP”), sacrificed, and their neural retina removed for mRNA extraction. The remaining mice were evaluated for bioluminescent signal before and after treatment with TMP by oral gavage (“+ TMP”), yielding the FLuc biolum. signal. After bioluminescence analysis, mice were also sacrificed and their neural retina removed for mRNA extraction. NLuc and FLuc were transcripts were evaluated and normalized to the RPLP2 housekeeping gene for each mouse.

**Sup. Fig. 3.**
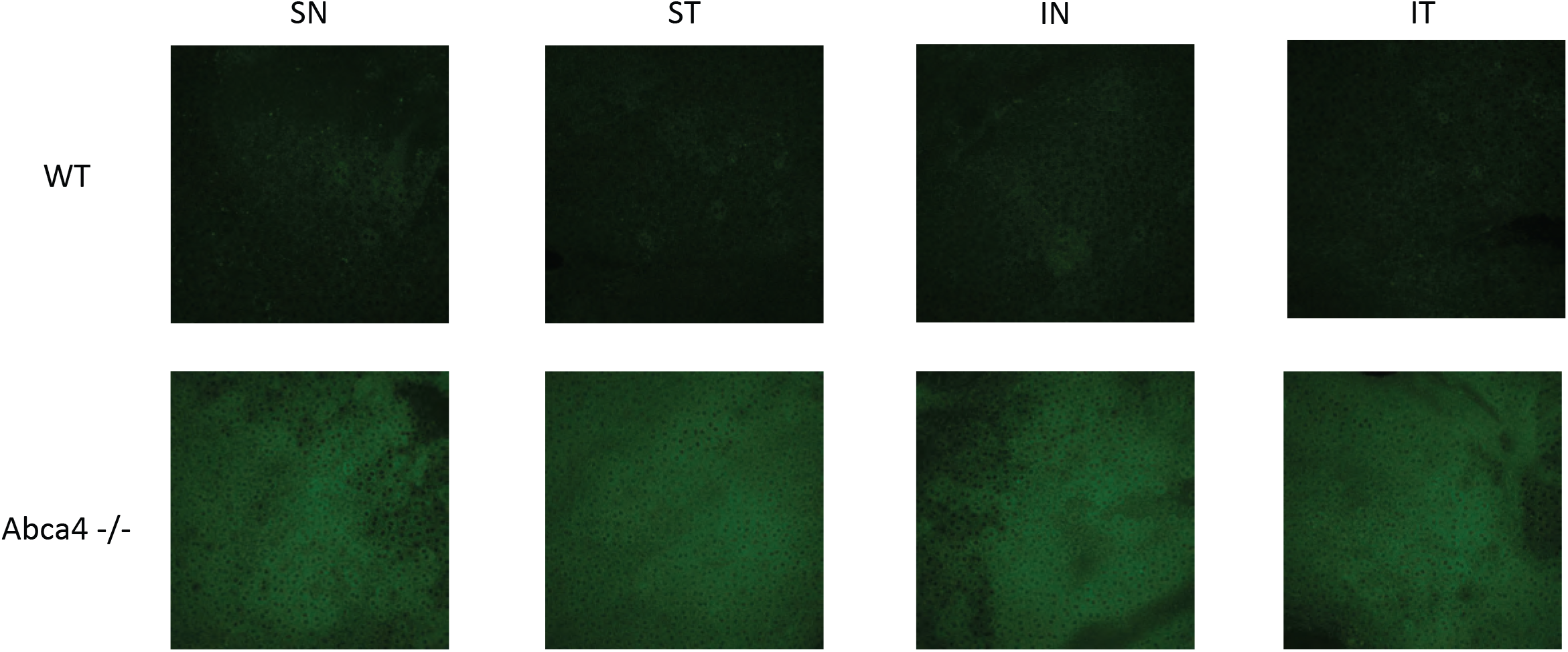
Confirmation of increased RPE autofluorescence in 6 mo *Abca4*^−/−^ mice. RPE flat mounts separated into quadrants were performed on WT and Abca4^−/−^ at 6 mo to confirm the Stargardt disease phenotype of increased levels of bisretinoid-mediated autofluorescence. SN – superior nasal, ST – superior temporal, IN – inferior nasal, IT – inferior temporal.

